# Discrete virus factories form in the cytoplasm of cells co-infected with two strains of the segmented dsRNA virus, infectious bursal disease virus (IBDV), that subsequently coalesce

**DOI:** 10.1101/870246

**Authors:** Elle A. Campbell, Alice G. Gray, Joanna Wells, Jennifer Simpson, Pippa C. Hawes, Andrew J. Broadbent

**Author notes:** Department of Biology & Biochemistry, University of Bath (Elle A. Campbell).

## Abstract

The *Birnaviridae* family, responsible for major economic losses to poultry and aquaculture, are non-enveloped viruses with a segmented double-stranded (ds)RNA genome that replicate in discrete cytoplasmic virus factories (VFs). Reassortment is common, however, the underlying mechanism remains unknown given that VFs may act as a barrier to genome mixing. In order to provide new information on VF trafficking during dsRNA virus co-infection, we rescued two recombinant infectious bursal disease viruses (IBDVs) of strain PBG98 containing either a split GFP11- or Tetracysteine (TC)- tag fused to the VP1 polymerase (PBG98-VP1-GFP11 and PBG98-VP1-TC). DF-1 cells transfected with GFP1-10 prior to PBG98-VP1-GFP11 infection, or stained with ReAsH following PBG98-VP1-TC infection, had green or red foci in the cytoplasm respectively that co-localised with VP3 and dsRNA, consistent with VFs. The average number of VFs decreased from a mean of 60 to 5 per cell between 10 and 24 hours post infection (hpi) (p<0.001), while the average area increased from 1.24 μm^2^ to 45.01μm^2^ (p<0.001), and live cell imaging revealed that the VFs were highly dynamic structures that coalesced in the cytoplasm. Small VFs moved faster than large (average 0.57μm/s at 16 hpi compared to 0.22 μm/s at 22 hpi), and VF coalescence was dependent on an intact microtubule network and actin cytoskeleton. During co-infection with PBG98-VP1-GFP11 and PBG98-VP1-TC viruses, discrete VFs initially formed from each input virus that subsequently coalesced 10-16 hours post-infection. We speculate that dsRNA virus reassortment requires VF coalescence, and the potential for reassortment increases at later time points in infection.

**Importance:** Reassortment is common in viruses with segmented double stranded (ds)RNA genomes. However, these viruses typically replicate within discrete cytoplasmic virus factories (VFs) that may represent a barrier to genome mixing. We generated the first replication competent tagged reporter birnaviruses, infectious bursal disease viruses (IBDVs) containing a split GFP11 or tetracysteine (TC) tag and used the viruses to track the location and movement of IBDV VFs, in order to better understand the intracellular dynamics of VFs from two different strains of dsRNA virus during a co-infection. Discrete VFs initially formed from each virus that subsequently coalesced from 10 hours pi. We hypothesise that VF coalescence is required for the reassortment of dsRNA viruses and the potential for reassortment increases later in the replication cycle. This study provides new information that adds to our understanding of dsRNA virus VF trafficking.

## Introduction

Members of the *Birnaviridae* family are responsible for some of the most economically devastating diseases to the poultry industry and aquaculture: Infectious bursal disease virus (IBDV) is endemic worldwide and ranks in the top 5 diseases of chickens in nearly all countries surveyed [1]. As well as causing severe morbidity and mortality, the virus is immunosuppressive, leaving birds that recover with an increased susceptibility to secondary infection and a reduced response to vaccination programs [2, 3]. Infectious Pancreatic Necrosis Virus (IPNV) is responsible for high mortalities in farmed salmon and trout and some strains can cause persistent infection, with fish spreading the virus by vertical or horizontal transmission [4]. In addition, more recently described birnaviruses, for example chicken proventriculus necrosis virus (CPNV) [5] and blotched snakehead virus (BSNV) [6] cause production loses that are only just beginning to be understood, and birnaviruses of insects such as Drosophila X virus (DXV) and Culex Y virus (CYV) are useful as tools for studying cellular antiviral responses [7]. However, despite the importance of these viruses, our understanding of how they replicate in cells is lacking.

The *Birnaviridae* genome is comprised of two segments of double stranded (ds)RNA. Segment A encodes two overlapping reading frames (ORFs), one encoding a non-structural protein (termed VP5 in IBDV and IPNV) and the other a polyprotein that is cleaved into the capsid protein (VP2), protease (VP4) and a dsRNA binding protein (VP3). Segment B contains one ORF encoding an RNA dependent RNA polymerase (VP1). Some VP1 copies bind the 5’ and 3’ ends of each genome segment and are packaged into the virion. The dsRNA genome is coated with VP3, which binds VP1 and activates its polymerase activity [8]. Collectively, VP1, VP3 and the dsRNA genome form a viral ribonucleoprotein (vRNP) complex [9].

The virus enters host cells by endocytosis or macropinocytosis. As the calcium concentration in the endosome drops, the virus uncoats, releasing a peptide which leads to the formation of pores in the endosomal membrane [10]. It is thought that the vRNP complexes exit the endosome through the pores to initiate transcription and translation and form a replication complex, or virus factory (VF) by co-opting endosomal membrane components [11, 12]. Unlike *Reoviridae*, which are also non-enveloped viruses with a segmented dsRNA genome, the *Birnaviridae* possess only one capsid and lack a transcriptionally active T2 core. It has been demonstrated that genome replication does not require the presence of the capsid [9, 13], and it is thought that the proteins in the vRNP complex shield the dsRNA genome from the cellular sensing machinery [14].

Little is known regarding the number, location and dynamics of birnavirus VFs during an infection. In contrast, the VFs of mammalian orthoreovirus are known to be highly dynamic, moving in the cytoplasm in a microtubule-dependent manner [15]. This is, in part, due to a lack of tagged reporter birnaviruses. Recently, a split GFP system has been used to generate tagged reporter viruses [16–19]. Briefly, the nucleotide sequence encoding the 11^th^ beta sheet of the GFP molecule (GFP11) is incorporated into the viral genome such that a viral protein tagged with the GFP11 peptide is translated. A plasmid encoding the rest of the protein (GFP1-10) is transfected into cells, and when the two come together, they produce a complete, fluorescing, GFP protein. A tetracysteine (TC) tag has also been used to engineer tagged reporter viruses [15, 20–26]. Briefly, the TC tag is comprised of 6 amino acids, including two separate pairs of cysteine residues (CCPGCC). When live infected cells are stained with biarsenical compounds (for example FLAsH-EDT_2_ or ReAsH-EDT_2_) the compounds covalently bind to the cysteine residues in the TC tag. This interaction leads to FLAsH-EDT_2_ fluorescing green and ReAsH-EDT_2_ fluorescing red [27]. Here, we describe the generation of the first ever replication competent tagged reporter birnaviruses, IBDVs tagged with either GFP11 or TC, and we use the viruses to describe the location and movement of *Birnaviridae* VFs in the cytoplasm.

As the *Birnaviridae* genome is divided into two segments, reassortment is a problem in the field. Reassortment of IBDV has been observed to occur between field strains and vaccine strains [28], between different serotypes [29], and is thought to be responsible for the emergence of very virulent strains [30], and reassortment has also been observed during IPNV infections [4, 31], complicating the epidemiology of the diseases. However, despite the importance of reassortment, the molecular mechanisms involved remain unknown. In order for reassortment to occur, the same cell must become co-infected and the viral gene segments must reach the same intracellular compartment. As the *Birnaviridae* replicate within discrete VFs, which may be a barrier for genome mixing, it remains unknown how genome segments reach the same intracellular compartment. Here, we track the location and movement of *Birnaviridae* VFs during co-infection of DF-1 cells with GFP11- and TC-tagged IBDVs in order to better understand the intracellular compartmentalisation of VFs from two different strains of dsRNA virus during a co-infection and therefore the potential for reassortment throughout the replication cycle.

## Results

### Construction of tagged reporter IBDV strains

A reverse genetics system developed in our laboratory was used to rescue a cell-culture adapted IBDV strain, PBG98. The PBG98 sequence was used as a backbone, and the nucleotide sequences encoding either the GFP11 or TC tags were added to the 3’ end of the coding region of Segment B, which encodes the VP1 polymerase (Fig. 1 A and B). Cells were transfected with the GFP1-10 molecule prior to PBG98-VP1-GFP11 infection, which allowed a full-length GFP protein to assemble in the cytoplasm, tagged to the C terminus of VP1. Multiple green foci were observed in the cytoplasm of infected cells, in contrast to cells transfected with GFP1-10 or GFP11 alone, which showed no positive signal, or cells transfected with GFP1-10 and subsequently transfected with a plasmid encoding GFP11, where a positive signal was observed throughout the cell (Fig. 1C). To visualise the TC tag, cells infected with the PBG98-VP1-TC virus were stained with ReAsH. Multiple red foci were observed in the cytoplasm of infected cells, in contrast to mock-infected cells also treated with ReAsH (Fig. 1D). The replication of both tagged viruses in DF-1 cells was compared to the recombinant wild-type (wt) PBG98 virus. In one experiment, the PBG98 virus replicated to a peak titre of 11.3 Log_10_TCID_50_/mL at 72hpi, whereas the PBG98-VP1-GFP11 virus only reached a titre of 8.9 Log_10_TCID_50_/mL (p<0.001) (Fig. 1E). In another experiment, the PBG98 virus replicated to a titre of 10.8 Log_10_TCID_50_/mL at 72hpi that was matched by the PBG98-VP1-TC virus, although replication was lower at earlier time points (p<0.05) (Fig. 1F). Taken together, both tagged viruses were attenuated, but the TC-tagged virus was less attenuated than the GFP11-tagged virus. The stability of the tags was determined by serially passaging the tagged-viruses ten times in DF-1 cells and then imaging DF-1 cells infected with the supernatants from each passage by confocal microscopy. The PBG98-VP1-GFP11 virus was stable up to 7 passages, whereas the TC-tag continued to be expressed even at passage 10, demonstrating that the PBG98-VP1-TC strain was more stable than the PBG98-VP1-GFP11 virus (Fig. S1).

**Fig 1.**
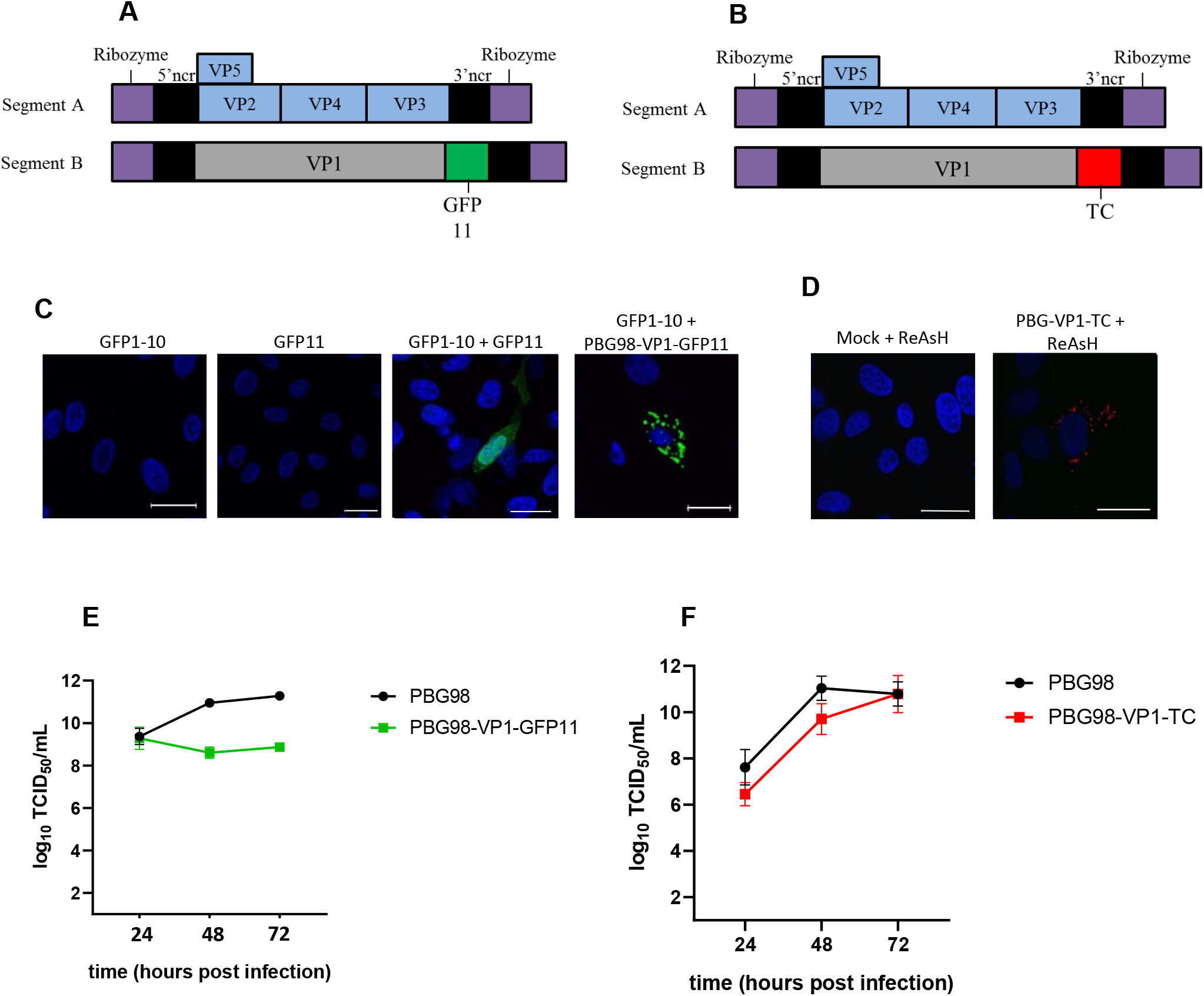
Construction of tagged reporter IBDV strains. The PBG98 sequence was used as a backbone, and the nucleotide sequences encoding either the GFP11-tag (A) or TC-tag (B) were added to the 3’ end of the coding region of Segment B. To obtain a positive GFP signal, DF-1 cells were transfected with a plasmid expressing GFP1-10, prior to transfection with a plasmid expressing GFP11, or infection with the PBG98-VP1-GFP11 virus (C). To visualise the TC tag, DF-1 cells infected with the PBG98-VP1-TC virus were stained with ReAsH (D). The replication of both tagged viruses in DF-1 cells was compared to the recombinant wild-type (wt) PBG98 virus. The titre to which the GFP11-tagged virus (E) or TC-tagged virus (F) replicated was compared to the wt PBG98 virus at the indicated time points post-infection in triplicate. Virus titres were expressed as Log_10_ TCID_50_/mL and the means plotted. Error bars represent the standard error of the mean (SEM) and the data are representative of 3 independent experiments (F).

### The PBG98-VP1-GFP signal is a marker for IBDV VFs

DF1 cells transfected with GFP1-10 and infected with the PBG98-VP1-GFP11 virus were fixed at 20 hpi and stained with either anti-dsRNA or anti-VP3 mouse monoclonal antibodies followed by a goat anti-mouse secondary antibody conjugated to Alexa Flour 568. The GFP11 signal was highly co-localised with both VP3 and dsRNA (Fig. 2 A and B), for example 82.6+/−0.1% of the signal derived from PBG98-VP1-GFP11 co-localised with the VP3 signal. Taken together, these data demonstrate that the PBG98-VP1-GFP11 signal co-localised with other vRNP components, consistent with the fluorescent signal from the tagged viruses being a marker for VFs.

**Fig. 2.**
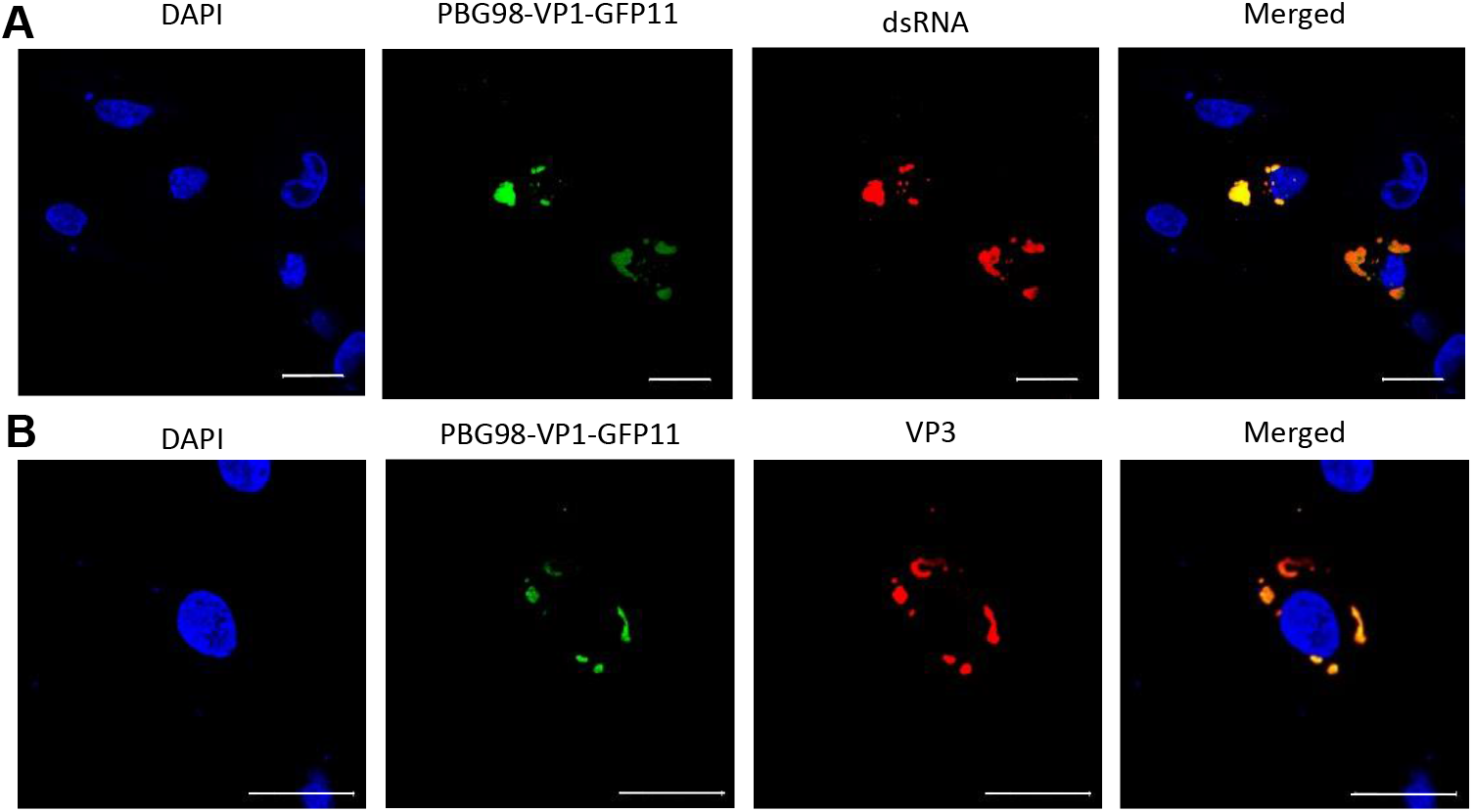
The PBG98-VP1-GFP signal is a marker for IBDV VFs. DF1 cells were transfected with a plasmid expressing GFP1-10 and infected 24 hours post-transfection with the PBG98-VP1-GFP11 virus at an MOI of 1. Cells were fixed at 20 hours post-infection and stained with DAPI and either anti-dsRNA (A) or anti-VP3 (B) mouse monoclonal antibodies followed by a goat anti-mouse secondary antibody conjugated to Alexa Flour 568, and imaged. Scale bars are 20μm in length.

### VFs coalesce in the cytoplasm of infected cells throughout infection

DF-1 cells were transfected with GFP1-10 and infected with the PBG98-VP1-GFP11 virus at an MOI of 1 and fixed at 10, 18 and 24 hpi. At early time points, distinct, small foci were abundant throughout the cytoplasm of infected cells, however, at later time points the foci were larger in size and fewer in number (Fig. 3A), To quantify this observation, 30 infected cells were imaged per time point. The mean number of foci significantly decreased from 60 per cell at 10 hpi to 5 per cell at 24 hpi (p< 0.001) (Fig. 3B) whereas the average area of each focus significantly increased from a mean of 1.2μm^2^ at 10 hpi to 45.0μm^2^ at 24 hpi (p<0.001) (Fig. 3C). Taken together, these data suggest that *Birnaviridae* VFs coalesced in the cytoplasm throughout infection. As multiple VFs were observed in infected cells, we sought to determine whether this was a feature of multiplicity of infection (MOI). To this end, we infected DF-1 cells with the PBG98-VP1-GFP11 virus at an MOI of 0.1, 0.01 and 0.001 and fixed and imaged cells 18 hours pi. Multiple VFs were observed in infected cells, irrespective of MOI (Fig. S2), indicating that this phenomenon was observed even in cells infected with a low amount of infectious virus.

**Fig. 3.**
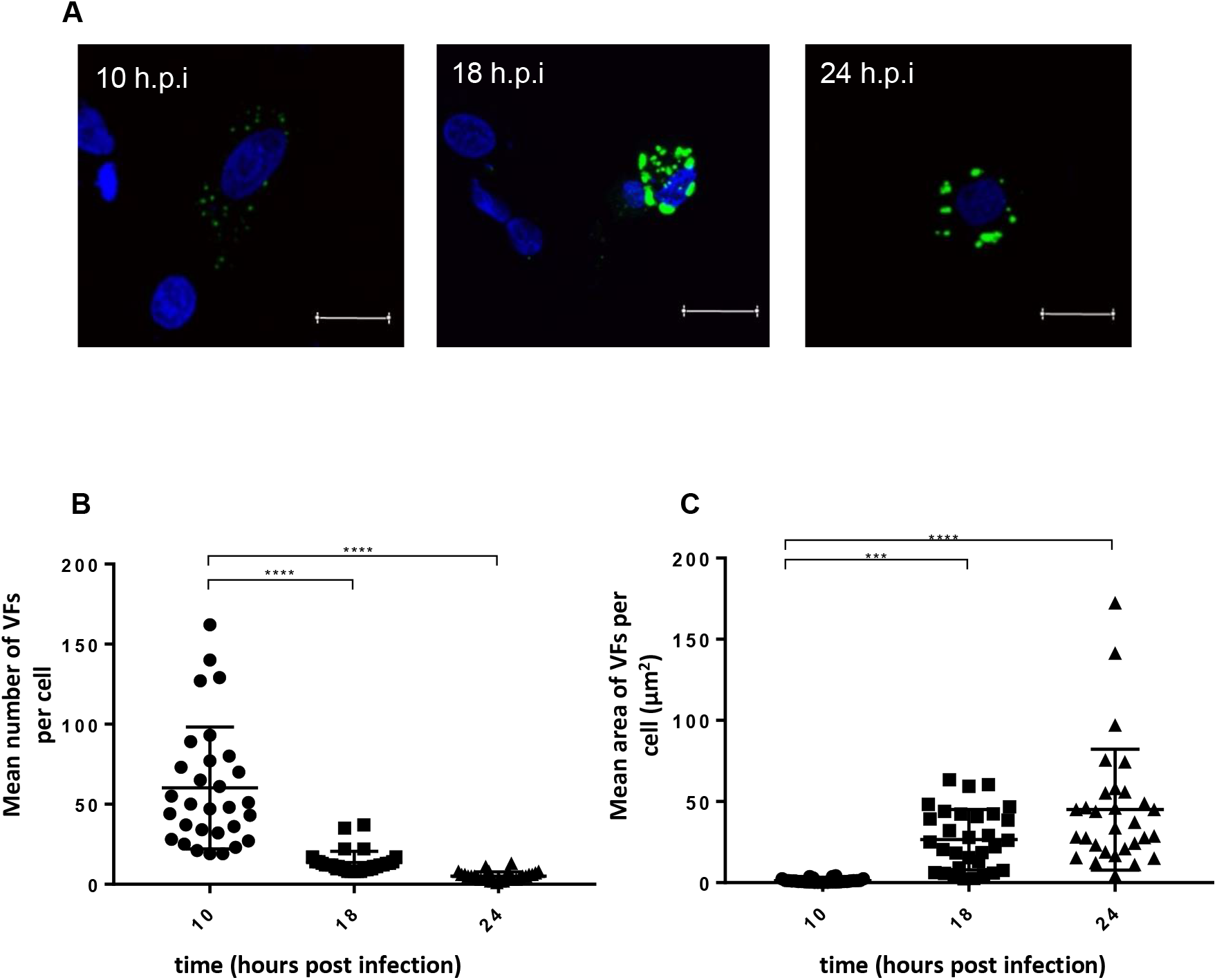
VFs decrease in number and increase in size over time. DF1 cells were transfected with a plasmid expressing GFP1-10 and infected 24 hours post-transfection with the PBG98-VP1-GFP11 virus at an MOI of 1. Cells were fixed at 10, 18, and 24 hours post-infection, stained with DAPI, and imaged (A). Scale bars are 20μm in length. The number of VFs per cell was determined for 30 infected cells at each time point and plotted (B). The average area of the VFs in an infected cell was determined using the surface tool in Imaris 9 software (bitplane) and plotted for 30 infected cells at each time point (C). The line represents the mean and the error bars represent the standard deviation (SD) of the mean.

Infected cells were subject to live cell imaging to investigate the nature of VF movement. Cells were imaged every 4 minutes over a 2 hour period from 16-18 hpi and a movie was made (Fig. 4A and Movie S1). Five fusion events were observed between foci in a single cell, and an example is shown in Fig. 4A. Fusion events were also apparent in cells imaged 22-24 hpi (Fig. 4B and Movie S2), although they were rarer at later time-points and in some cases transient. Smaller foci were also observed splitting away from larger foci. In addition, there was a significant difference in speed of foci movement between 16 and 22 hpi. At 16 hpi, foci moved with an average speed of 0.57μm/s, while at 22 hpi, foci only moved with an average speed of 0.22μm/s (p<0.001). Taken together, these data demonstrate that *Birnaviridae* VFs are dynamic structures that coalesce in the cytoplasm.

**Fig. 4.**
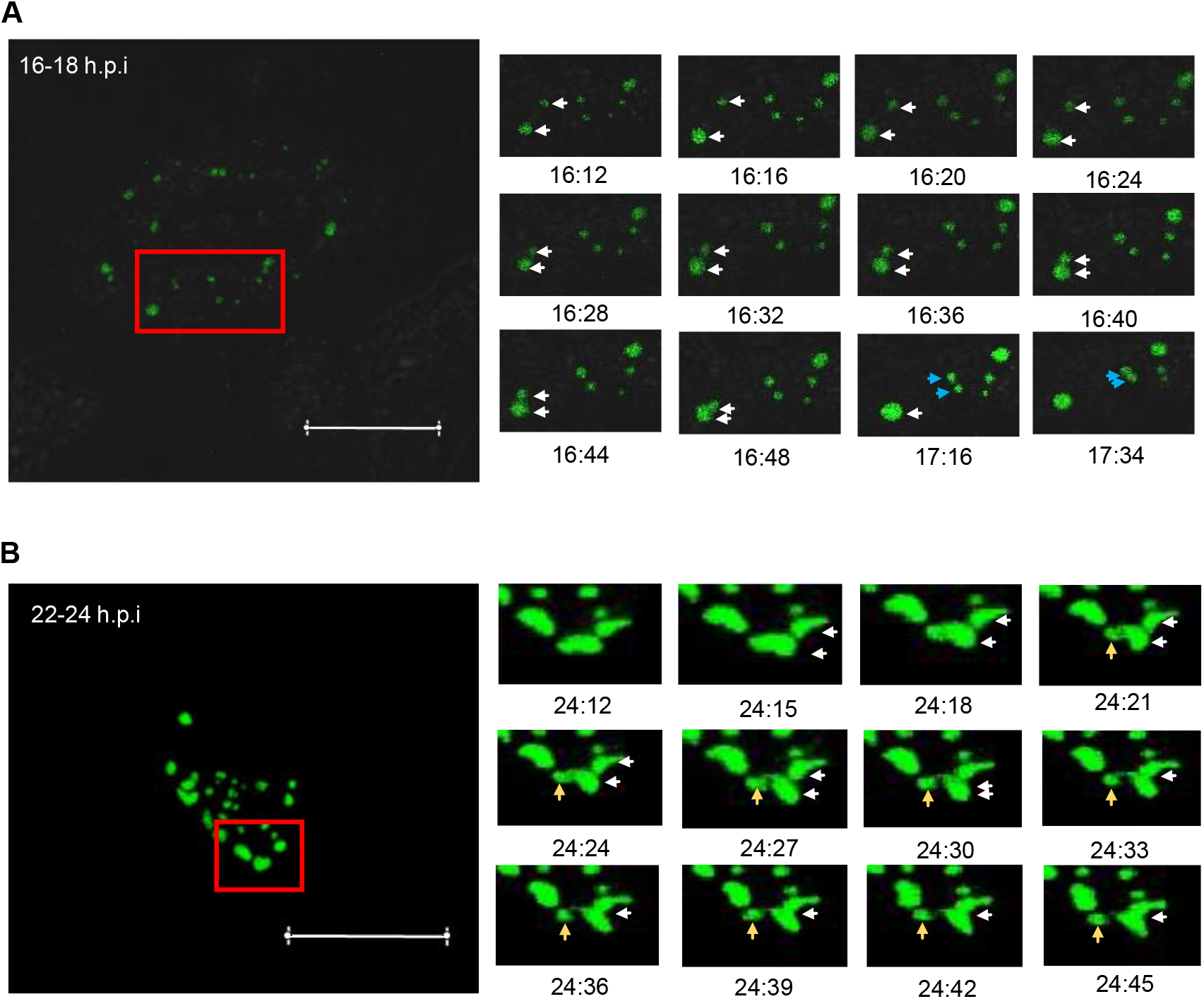
VFs coalesce in the cytoplasm of infected cells throughout infection. DF1 cells were transfected with a plasmid expressing GFP1-10 and infected 24 hours post-transfection with the PBG98-VP1-GFP11 virus at an MOI of 1. One live infected cell was imaged from 16-18 hours post infection. Images of the boxed region are shown at the indicated time points post infection where two VF coalescence events were witnessed (white arrows and blue arrows) (A). Another live infected cell was imaged from 22-24 hours post-infection. Images of the boxed region are shown at the indicated time points post infection where one VF coalescence event was witnessed (white arrows) and one VF splitting event was witnessed (yellow arrows) (B).

### VF coalescence is dependent on an intact microtubule network and actin cytoskeleton

Cells were infected with the PBG98-VP1-TC virus and then stained with ReAsH and fixed and stained with anti-tubulin and a secondary antibody conjugated to Alexa Flour 488 (Fig. 5A). While little co-localisation was observed between the VFs and the microtubule network, when cells were infected with the PBG98-VP1-TC virus and then stained with ReAsH and Phalloidin -Alexa Fluor 488, the edges of the red foci co-localised with the Phalloidin (Fig. 5B), suggesting that actin filaments may be involved in VF movement. To determine whether the microtubule network or actin cytoskeleton were involved in *Birnaviridae* VF coalescence, cells were treated with either nocodazole, which triggers microtubule depolymerisation, or cytochalasin D, which inhibits actin filament polymerisation, 2 hours following infection with the PBG98-VP1-TC virus. Actin is known to play a role in IBDV entry [32] and so this time-point was selected to distinguish the effect of drug treatment on VF movement from virus entry. At 18 hpi, both nocodazole and cytochalasin D treated cells had significantly more foci per cell (a mean of 25 and 30 foci per cell, respectively) than mock-treated infected cells (a mean of 7 foci per cell), p<0.0001 (Fig. 5C). Nocodazole and cytochalasin D treatments also led to a significantly reduced area of foci at this time point (14.4μm^2^ and 8.1μm^2^, respectively) compared to mock-treated infected controls (32.1μm^2^), p<0.0001 (Fig. 5D). Despite the differences in VF size and number following treatment, neither drug caused significant differences in the titre of the PBG98-VP1-TC virus recovered from the supernatant of infected cultures at 24 hpi (Fig. 5E). Cells were transfected with GFP1-10 and infected with PBG98-VP1-GFP11 at an MOI of 1, treated with either nocodazole or cytochalasin D at 2 hpi, and imaged live from 20-22 hpi. In cells treated with nocodazole, multiple groups of small foci were seen to go through series of splitting and fusion events, at speeds comparable to when no drug treatment was applied (0.62μm/s), but the small foci did not coalesce into larger foci over this time period (Fig. 6A and Movie S3). In contrast, while foci continued to move in cells treated with cytochalasin D, there was no visual sign of any foci interacting over the time of imaging, and the speed of movement was significantly reduced compared to mock-treated infected cells, with foci moving with a mean speed of only 0.17μm/s (p<0.0001) (Fig. 6B and Movie S4). Taken together, these data demonstrate that birnavirus VF coalescence is dependent on both an intact microtubule network and actin cytoskeleton and that inhibition of actin polymerisation significantly slows birnavirus VF movement.

**Fig. 5.**
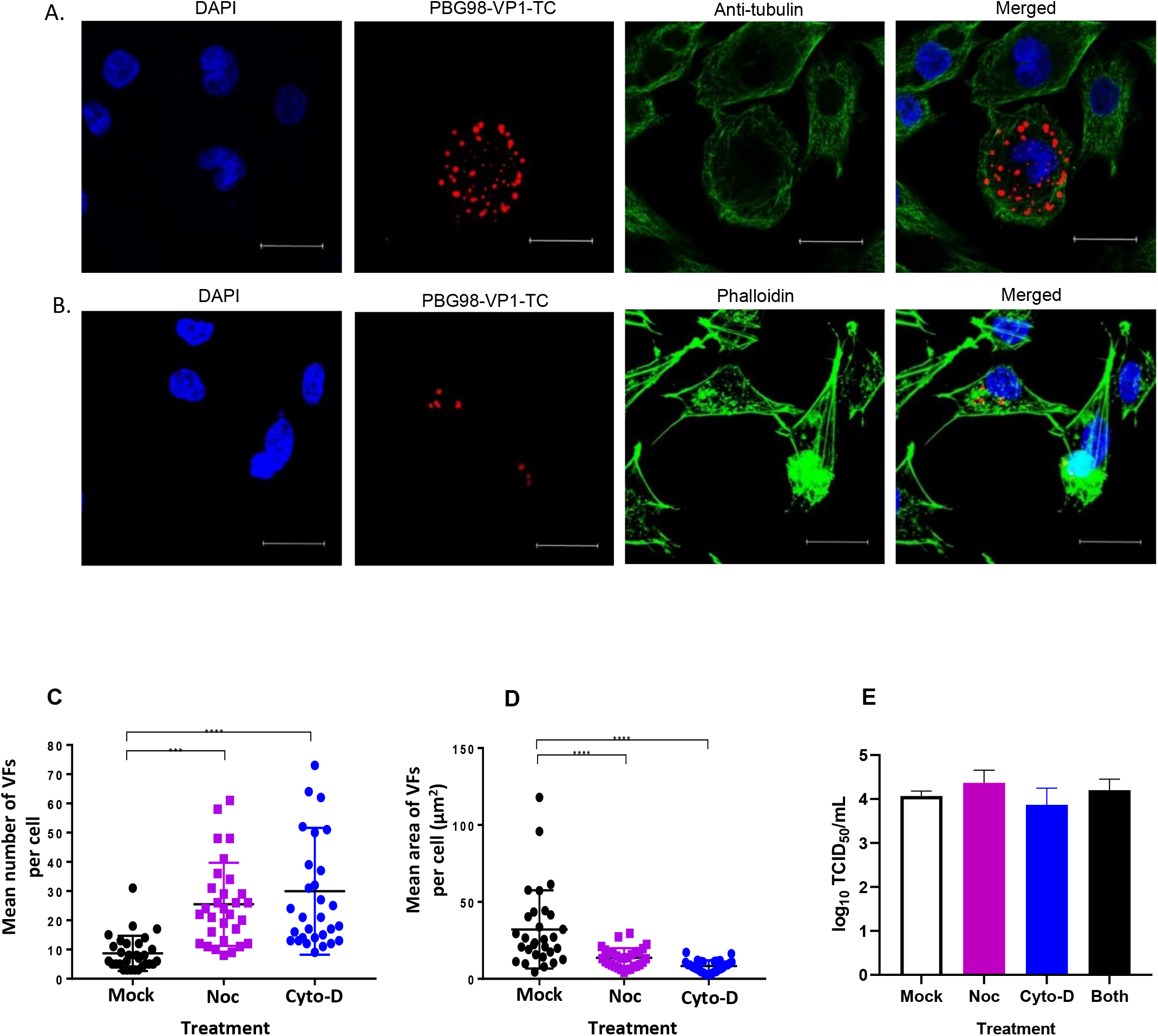
The distribution of VFs in the cytoplasm is dependent on an intact actin cytoskeleton and microtubule network, but alterations in VF distribution do not alter viral replication. DF1 cells were infected with the PBG98-VP1-TC virus at an MOI of 1. Cells were fixed at 20 hours post-infection and stained with DAPI and either an anti-tubulin mouse monoclonal antibody followed by a goat anti-mouse secondary antibody conjugated to Alexa Flour 488 to visualise the microtubule network (A) or phalloidin conjugated to Alexa Flour 488 to visualise the actin cytoskeleton (B). Scale bars are 20μm in length. The number of VFs per cell was determined for 30 infected cells and plotted (C). The average area of the VFs in an infected cell was determined using the surface tool in Imaris 9 software (bitplane) and plotted for 30 infected cells (D). The line represents the mean and the error bars represent the standard deviation (SD) of the mean. Cell supernatants obtained 24 hours post-infection were titrated and the tissue culture infectious dose 50 (TCID_50_) determined (E).

**Fig. 6.**
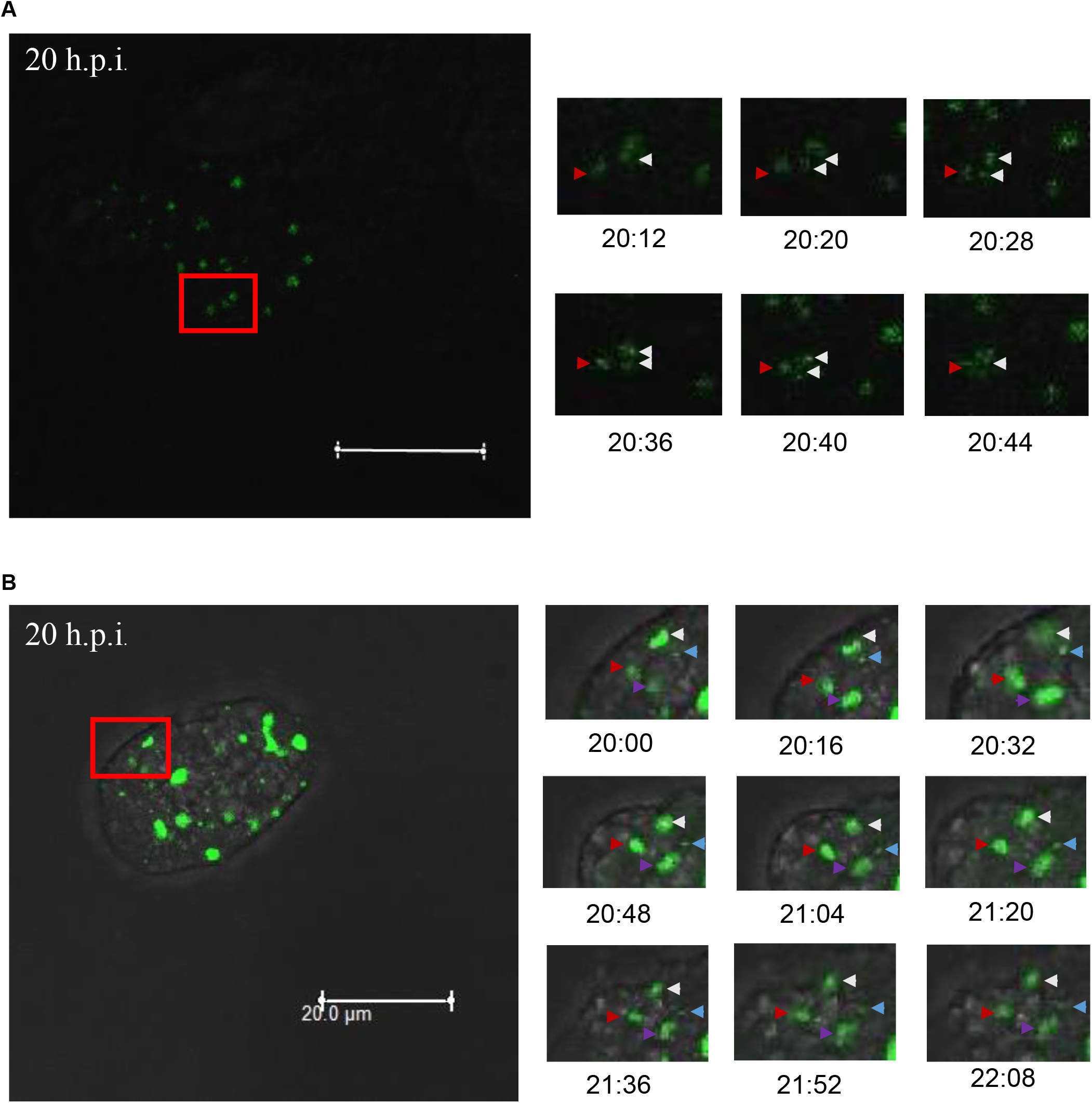
VF coalescence is dependent on an intact microtubule network and actin cytoskeleton. DF1 cells were transfected with a plasmid expressing GFP1-10 and infected 24 hours post-transfection with the PBG98-VP1-GFP11 virus at an MOI of 1. Two hours post-infection, cells were treated with either nocodazole (A) or cytochalasin-D (B). One live infected cell was imaged from 20-22 hours post infection. Images of the boxed region are shown at the indicated time points post infection. In an infected cell treated with nocodazole, one VF was witnessed splitting into two VFs that then subsequently coalesced (white arrows) next to a neighbouring VF (red arrows) (A). In an infected cell treated with cytochalasin-D, four VFs were observed (white, blue, red and purple arrows) that did not coalescence or interact with each other over the 2 hour period (B). Scale bars are 20μm in length.

### Discrete VFs from different strains of IBDV form in the cytoplasm of co-infected cells and coalesce throughout infection

Cells were transfected with GFP1-10 and co-infected with both the PBG98-VP1-GFP11 and PBG98-VP1-TC viruses at an MOI of 1, and fixed, ReAsH and DAPI stained at 10, 14, 18 and 24 hpi. Like in previous experiments, foci at later time points were less numerous and larger than foci at earlier time points, likely as a result of VF coalescence throughout infection (Fig. 7A). At 10 hpi, distinct green and red foci were observed in the cytoplasm of co-infected cells, with little evidence of co-localisation, consistent with the presence of discrete VFs from the PBG98-VP1-GFP11 and PBG98-VP1-TC viruses in the cytoplasm. However, at 14, 18 and 24 hpi, there was considerable co-localisation between red and green foci (Fig. 7A), consistent with the coalescence of VFs from the PBG98-VP1-GFP11 and PBG98-VP1-TC viruses. Data derived from Image J’s co-localisation function demonstrated that the levels of co-localisation between green and red foci increased from 18.2% at 10 hpi to 70.6% at 24 hpi (p<0.0001) (Fig. 7B). To better define the timing of coalescence between VFs, DF-1 cells were transfected with GFP1-10 and co-infected with both the PBG98-VP1-GFP11 and PBG98-VP1-TC viruses at an MOI of 1, and fixed, ReAsH, and DAPI stained at 10, 12, 14 and 16 hours post infection. Thirty co-infected cells were imaged per time-point and the average percentage of discrete red and green foci, and co-localised (yellow) foci quantified per cell. At 10 hpi, no co-localisation was observed between red and green foci in co-infected cells. However, an average of 13.1% of red and green foci within co-infected cells co-localised with each other at 12 hpi. This increased to 36.5% and 89.9% at 14 and 16 hpi (Fig. 7C, yellow bars) consistent with the coalescence of VFs from the PBG98-VP1-GFP11 and PBG98-VP1-TC viruses between 10 and 16 hours pi. Moreover, cytochalasin D treatment delayed the coalescence of red and green VFs as the average percentage of co-localised red and green foci per co-infected cell was only 0.9% and 18.8% at 12 and 14 hpi in the presence of the drug (Fig. 7C). Taken together, these data demonstrate that during co-infection, discrete *Birnaviridae* VFs form in the cytoplasm from each input virus that subsequently coalesce over time.

**Fig. 7.**
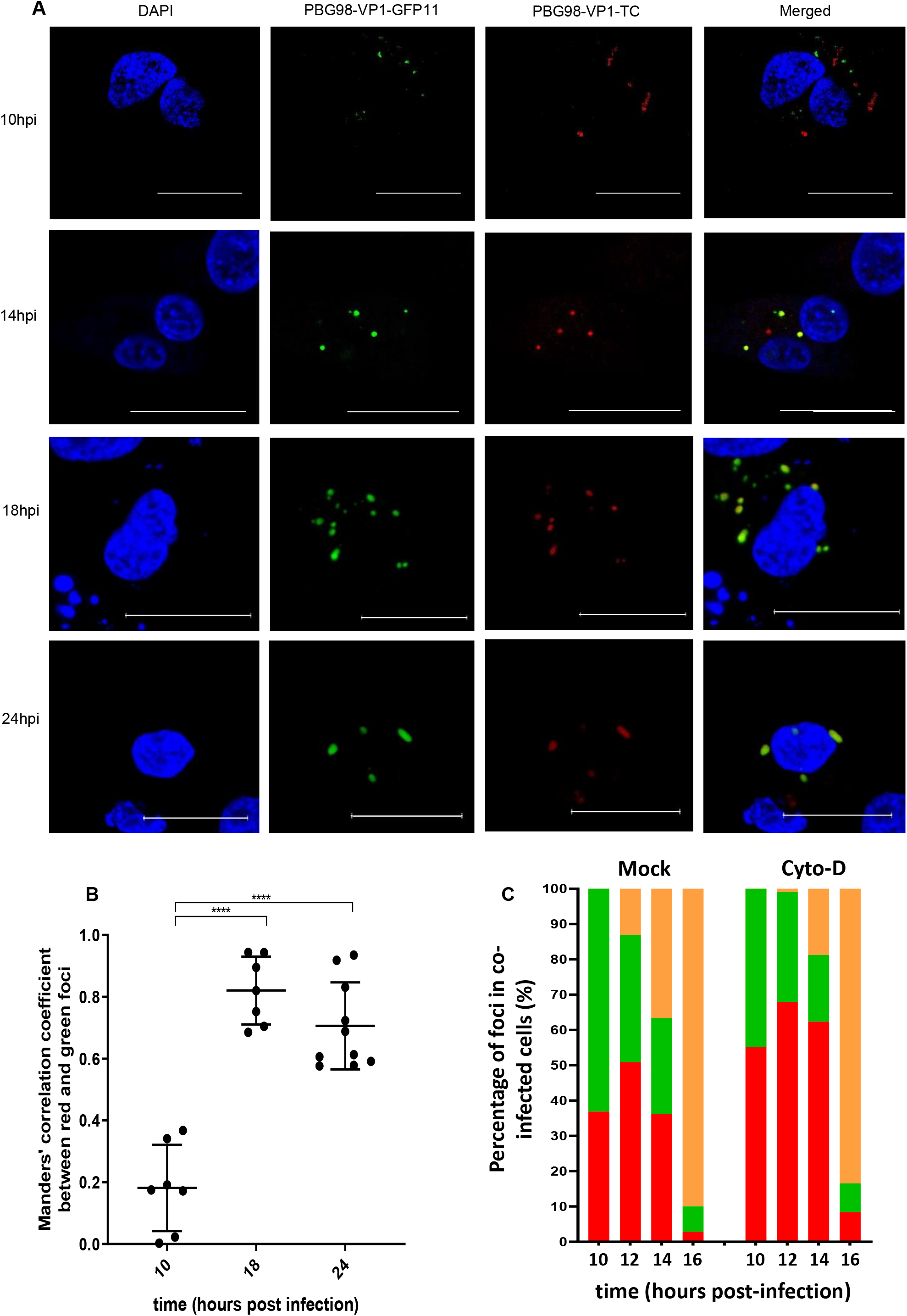
Discrete VFs from different strains of IBDV form in the cytoplasm of co-infected cells and coalesce throughout infection. DF1 cells were transfected with a plasmid expressing GFP1-10 and co-infected 24 hours post-transfection with the PBG98-VP1-GFP11 virus and the PBG98-VP1-TC virus at an MOI of 1. Cells were fixed at 10, 14, 18 and 24 hours post-infection and stained with ReAsH and DAPI; Scale bars are 20μm in length (A). The Mander’s correlation coefficient between red and green foci was plotted for 10 cells at 10, 18 and 24 hours post-infection. The line represents the mean and the error bars represent the standard deviation (SD) of the mean (B). The percentage of red, green, and co-localised (yellow) foci were quantified for 30 co-infected cells per time point and the average plotted at 10, 12, 14 and 16 hours post infection (C).

## Discussion

In this study, we produced the first replication competent tagged-reporter birnaviruses, an IBDV tagged with GFP11 (PBG98-VP1-GFP11) and an IBDV tagged with TC (PBG98-VP1-TC). We used the PBG98-VP1-GFP11 virus to directly visualise IBDV VFs in live infected cells and demonstrated that the VFs were dynamic structures that moved within the cytoplasm and coalesced during the course of infection. These findings are in agreement with others who have found IBDV replication complexes to be located in distinct cytoplasmic puncta [11] that increase in size over the course of infection [13] by imaging fixed cells, however, our data extend these results by imaging live infected cells and characterising VF movement. The VFs of mammalian orthoreovirus (MRV) were also recently found to be highly dynamic structures that move and interact in the cytoplasm [15], increasing in size and decreasing in number, consistent with coalescence [33]. Therefore, it is possible that the dynamic nature and coalescence of VFs may be a common feature of dsRNA viruses.

We found that destabilisation of the microtubule network with nocodazole had a significant impact on IBDV VF size, number, and dynamics throughout infection (Fig. 5 and 6A and Movie S3). Consistent with this observation, Delgui et al found that when IBDV-infected quail muscle (QM7) cells were treated with nocodazole, more VFs were situated at the periphery of the cytoplasm, failing to traffic to the perinuclear region, unlike in mock-treated cells [11]. The distribution of IBDV VFs in our study was less restricted to the perinuclear region in the absence of drug treatment, however, we used DF-1 cells in our experiments which may account for this discrepancy. Moreover, the distribution of VFs we saw is consistent with data from Dalton & Rodriguez who also used DF-1 cells [13]. Interestingly, *Reoviridae* VFs are also known to hijack the microtubule network during intracellular trafficking [33], and utilisation of this cytoskeletal component may be a common feature among dsRNA virus families. We also found that actin filaments co-localised with the periphery of PBG98-VP1-TC VFs and that VF size and speed of movement and fusion events were significantly reduced following cytochalasin D treatment, suggesting that the actin cytoskeleton is also involved with *Birnaviridae* VF trafficking (Fig. 5 and 6B and Movie S4). Interestingly, VFs have also been observed co-localising with actin in cells infected with Fowlpox [34] and Bunyamwera virus [35], and actin is involved in the internalisation, replication, and non-lytic egress of rotavirus [36, 37]. IBDV internalisation into the cell is also dependent on an intact actin network [32], [38]. In order to distinguish between the involvement of actin in IBDV entry and VF movement, cultures were treated with cytochalasin D from 2 hours pi, to allow IBDV time to enter the cells prior to treatment. Drug treatment did not significantly alter virus replication (Fig. 5E) demonstrating that the replication of IBDV is not affected by alterations to VF trafficking, consistent with observations made by Delgui et al [11].

While the dynamics of VF movement has been previously demonstrated for other dsRNA viruses, our study is the first to image dsRNA VFs in co-infected cells. We demonstrated that during co-infection with two IBDV strains, discrete VFs from each input virus initially formed in the cytoplasm that subsequently coalesced over time, with the proportion of VFs containing both GFP11- and TC- tagged VP1 proteins increasing throughout the replication cycle (Fig. 7). The diversity of RNA viruses is increased through recombination and/or reassortment in cells co-infected with multiple strains. In order for this to occur, the genome from two or more infecting viruses must reach the same intracellular compartment. Using fluorescently tagged vaccinia viruses, Paszkowski et al demonstrated that during co-infection, distinct VFs from two input viruses formed in the cytoplasm and subsequently merged. The authors concluded that the VFs acted as a barrier to genomic mixing and that the merger of VFs was necessary for recombination [39]. It has also been proposed that the coalescence of VFs must take place in order for reassortment of dsRNA viruses to occur [40], in contrast to viruses such as influenza, where genome segments coalesce en route to the plasma membrane [41] and where reassortment is not restricted by VF compartmentalisation. Our data provide the first experimental demonstration of VF coalescence for the *Birnaviridae* and, to our knowledge, the first evidence of coalescence between VFs of different strains in a co-infected cell for any dsRNA virus family. Moreover, our data suggest that the potential for IBDV reassortment occurs late in the viral replication cycle, as VF coalescence between GFP11- and TC- tagged viruses did not occur until after 10 hpi. This study therefore provides new information that contributes to our understanding of VF trafficking in dsRNA viruses, and the molecular basis of reassortment.

Our study is not without limitations; for example, the fluorescent signal was only detected from 8 hpi, and was only bright enough to be reliably imaged from 10 hpi for either the GFP11- or the TC-tagged viruses. At 10 hpi, numerous small VFs were observed (Fig 3A), even in cells infected with a very low MOI (Fig S2), implying that the VFs are not the product of one cell being infected with multiple infectious viruses. The molecular basis of this observation remains unknown, and, as we were unable to study events earlier than 8 hpi, this remains beyond the scope of the current study.

We were also unable to rescue a replication competent PBG98-VP1-GFP virus. However, GFP11 and GFP1-10 come together to give a fluorescent signal, indicating that the VP1-GFP molecule is functional. The crystal structure of the birnavirus VP1 has been determined [42, 43] and our data demonstrate that the C terminal extension is able to tolerate the insertion of a tag, without blocking the function of the active site. However, the tagged viruses were both attenuated compared to the recombinant wt PBG98 strain (Fig. 1E and F). This could be because the presence of the tag negatively affected the function of the VP1 protein, or because the presence of the nucleotide sequence that encoded the tag made the genome segment less efficient at being packaged. Moreover, the GFP11-tagged virus was more attenuated and less stable than the TC-tagged virus (Fig.1 and S1), which correlates with the length of the tag. Investigating the reason for the differences in attenuation was beyond the scope of this project. In addition, some VP1 is packaged into the virion, while the rest remains in the cytoplasm, and some VP1 is bound to the dsRNA genome, while the rest remains unbound. It is unknown whether the VP1-GFP is packaged into the virion, or whether it is bound to the genome. Discerning these differences was beyond the scope of this project. As the TC tag was less attenuated, it can be considered as an accurate representation of IBDV-derived VFs in infected cells. However, the fluorescence produced by the staining of the TC tag with ReAsH was more prone to bleaching than the GFP signal, so the GFP11-tagged IBDV strain was a more attractive candidate for live cell imaging over long time courses. In order to detect a green signal, it was necessary for cells to be both successfully transfected with GFP1-10 and infected with the GFP11-tagged virus. While this approach is adequate for imaging experiments, we are currently establishing a DF1 cell –line that stably expresses the GFP1-10 molecule for additional studies. Finally, efforts to rescue an IBDV with a tag on the C terminus of Segment A were unprofitable. Segment A is translated as a polyprotein encoding VP2-VP4-VP3 which is subsequently cleaved [44]. The C terminus of VP3 has recently been shown to be essential for function [45], which might explain why this was an unsuitable position for the tag, however given that there are cleavage sites at the VP2-VP4 and VP4-VP3 junctions, it may be difficult to generate a tagged segment A.

Taken together, our data demonstrate that the *Birnaviridae* VFs are highly dynamic structures, moving and coalescing in the cytoplasm and that during co-infection with two strains of IBDV, discrete VFs form from each virus that subsequently coalesce. This study provides new information that adds to our understanding VF trafficking that could have implications for the molecular basis of dsRNA virus reassortment.

## Materials and Methods

### Cell-lines and antibodies

DF1 cells (chicken embryonic fibroblast cells, ATCC number CRL-12203) were sustained in Dulbecco’s modified Eagle’s medium (DMEM) (Sigma-Aldrich, Merck, UK) supplemented with 10% heat inactivated foetal bovine serum (hiFBS) (Gibco, ThermoFisher Scientific, UK). The primary antibodies used in this study were raised against tubulin (SantaCruz Biotechnology, UK), dsRNA (English & Scientific Consulting Kft.), and VP3 [46]. In all immunofluorescent assays, primary antibodies were diluted 1:100 and secondary antibodies conjugated to Alexa 488-or 568 (Invitrogen, Thermo Fisher Scientific, UK) were diluted 1:500 in a solution of BSA (Sigma-Aldrich, Merck, UK).

### Plasmids and recombinant viruses

The sequence of segments A and B of the cell-culture adapted IBDV strain, PBG98, including the 5’ and 3’ non coding regions (ncr’s) were flanked by self-cleaving ribozymes (a hammerhead ribozyme upstream and a hepatitis delta ribozyme downstream) and the whole sequence was ordered (GeneArt, ThermoFisher Scientific, UK) and cloned into a pSF-CAG-KAN vector (Sigma-Aldrich, Merck, UK) to make two “reverse genetics” plasmids (pRGs), pRG-PBG98-A and pRG-PBG98-B. DF-1 cells at 70% confluency were transfected with both plasmids with lipofectamine 2000 (Invitrogen, ThermoFisher scientific, UK) in order to rescue the recombinant PBG98 virus. Three alanine residues were added to the N terminus of the GFP11 tag as a linker and a stop codon was added to the C terminus to make the amino acid sequence: AAARDHMVLHEYVNAAGIT-Stop. The nucleotide sequence encoding this was added to the 3’ end of the coding region of segment B, which encodes VP1, prior to the 3’ ncr, to make the plasmid pRG-PBG98-B-GFP11 (GeneArt, ThermoFisher Scientific, UK). DF-1 cells were co-transfected with this plasmid and pRG-PBG98-A to generate the recombinant virus PBG98-VP1-GFP11. The tetracysteine tag (amino acid sequence: CCPGCC) was incorporated into the same region as the GFP11 tag to make the pRG-PBG98-B-TC plasmid (GeneArt, ThermoFisher Scientific, UK) and DF-1s were co-transfected with this plasmid and pRG-PBG98-A to make the recombinant virus PBG98-VP1-TC.

### Visualising virus infection

To visualise PBG98-VP1-GFP11 virus infection, DF1 cells were seeded onto coverslips (TAAB, UK) in 24-well plates (Falcon, Corning, UK) at a density of 1.6×10^5^ per well and transfected with GFP1-10 using lipofectamine 2000 24 hours prior to infection with the PBG98-VP1-GFP11 virus. Unless otherwise stated, cells were infected at an MOI of 1. To visualise PBG98-VP1-TC virus infection, live infected cells were stained with the TC-ReAsH™ II In-Cell Tetracysteine Tag Detection Kit (Invitrogen, ThermoFisher Scientific, UK), according to the manufacturer’s instructions.

### Immunofluorescence Microscopy

Cells were fixed with a 4% paraformaldehyde solution (Sigma-Aldrich, Merck, UK) for 20 minutes, permeabilized with a solution of 0.1% Triton X-100 (Sigma-Aldrich, Merck, UK) for 15 minutes and blocked with a 4% BSA solution for 30 minutes. Cells were then incubated with the appropriate primary antibody for one hour at room temperature. Cells were washed with PBS and incubated with the corresponding secondary antibody for 1 hour at room temperature in the dark. Cells were again washed and incubated in a solution of 4,6-diamidino-2-phenylindole dihdrochloride (DAPI, Invitrogen ThermoFisher Scientific, UK). Cells were washed in water, mounted with Vectashield (Vector Laboratories Inc, CA) and imaged with a Leica TCS SP5 confocal microscope. To image actin, Alexa Fluor™ 488 Phalloidin (Sigma-Aldrich, Merck, UK) was added directly to cells in PBS after blocking and incubated at room temperature for 25 minutes prior to DAPI staining.

### Live cell Imaging

DF1 cells were seeded into a Chambered #1.0 Borosilicate Coverglass slide (Nunc, Lab-Tek, Sigma-Aldrich, Merck, UK) at a density of 8 × 10^4^ per well and cultured in 1ml of DMEM supplemented with 10% hiFBS. Cells were transfected with GFP1-10 and then infected with the PBG98-VP1-GFP11 virus 24 hours post-transfection. Cells were maintained in Leibovitz’s L-15 media without phenol red (Gibco, ThermoFisher Scientific, UK) during live cell imaging experiments. Unless otherwise stated, 10-Z stacks were imaged every 4 minutes for a minimum of 2 hours using a Leica TCS SP8 confocal microscope.

### Imaging Quantification

The mean area and number of VFs were calculated using the surface tool in Imaris 9 software (Bitplane, Oxford Instruments, UK), and an overlap coefficient, (an alternative to the Pearson’s correlation coefficient created by Manders et al) was used as the main statistical test to describe co-localisation using Image J software (National Institutes of Health, NIH). Images were only considered in the analysis if they had a Coste’s significant level of 0.95 or above [17]. For live cell imaging, statistics such as displacement, velocity and number of merging and splitting events were calculated using the Image J plugin ‘TrackMate’.

### Virus Titration

Samples were titrated in 96-well plates (Falcon, Corning, UK) seeded with DF1 cells at a density of 4 × 10^4^ cells per well. Wells were then infected in quadruplicate with a 10-fold dilution series of viral supernatant. After 5 days, wells were inspected for signs of cytopathic effect (cpe) and the titre of virus determined by Tissue Culture Infective Dose-50 (TCID_50_) as per the method by Reed and Muench [47].

### Virus Growth curves

DF1 cells were seeded into 24 well plates at a density of 1.6 × 10^5^ cells per well in triplicate for each time point. The next day, cells were infected with either PBG98-VP1-GFP11, PBG98-VP1-TC or PBG98 viruses at an MOI of 0.01. Cell supernatant was then collected at 24, 48 and 72 hours post-infection and titrated as per the method by Reed and Muench [47].

### Passage Stability

DF1 cells were seeded into 24 well plates at a density of 1.6 × 10^5^ per well and maintained in 900μl media overnight. Cells were subsequently infected with 100μl of the supernatant collected from the previous passage. After 24 hours, the supernatant was collected and frozen at −80°C until the next passage. Both fluorescently tagged IBDV-viruses were passaged ten times, whereupon DF-1 cells were infected with the supernatant from every passage in order to image infected cells 20 hours post infection. Cells were transfected with GFP1-10 prior to infection with PBG98-VP1-GFP11 supernatants and were stained with ReAsH subsequent to infection with PBG98-VP1-TC supernatants.

### Drug treatment

DF1 cells were seeded onto coverslips and infected with the PBG98-VP1-TC virus. Cultures were subsequently treated with either cytochalasin D or nocodazole (Sigma-Aldrich, Merck, UK). Drugs were dissolved in Dimethyl sulfate (DMSO) (Sigma-Aldrich, Merck, UK) and diluted with media to a final concentration of 1 and 10 μM, respectively. Infected cultures were treated with drugs from 2 hours post infection until the end of the experiment. An equivalent volume of DMSO was added to the culture media of mock-treated control cells.

## Acknowledgements

We are grateful for financial support from the BBSRC (grants: BBS/E/I/00001845, BBS/E/I/00007034 and BBS/E/I/00007039). The funders had no role in study design, data collection and interpretation, or the decision to submit the work for publication. We acknowledge Dr Michael A Skinner, Imperial College London, for providing the sequence of PBG98 and the anti-VP3 antibody.

---

**Fig. S1.**
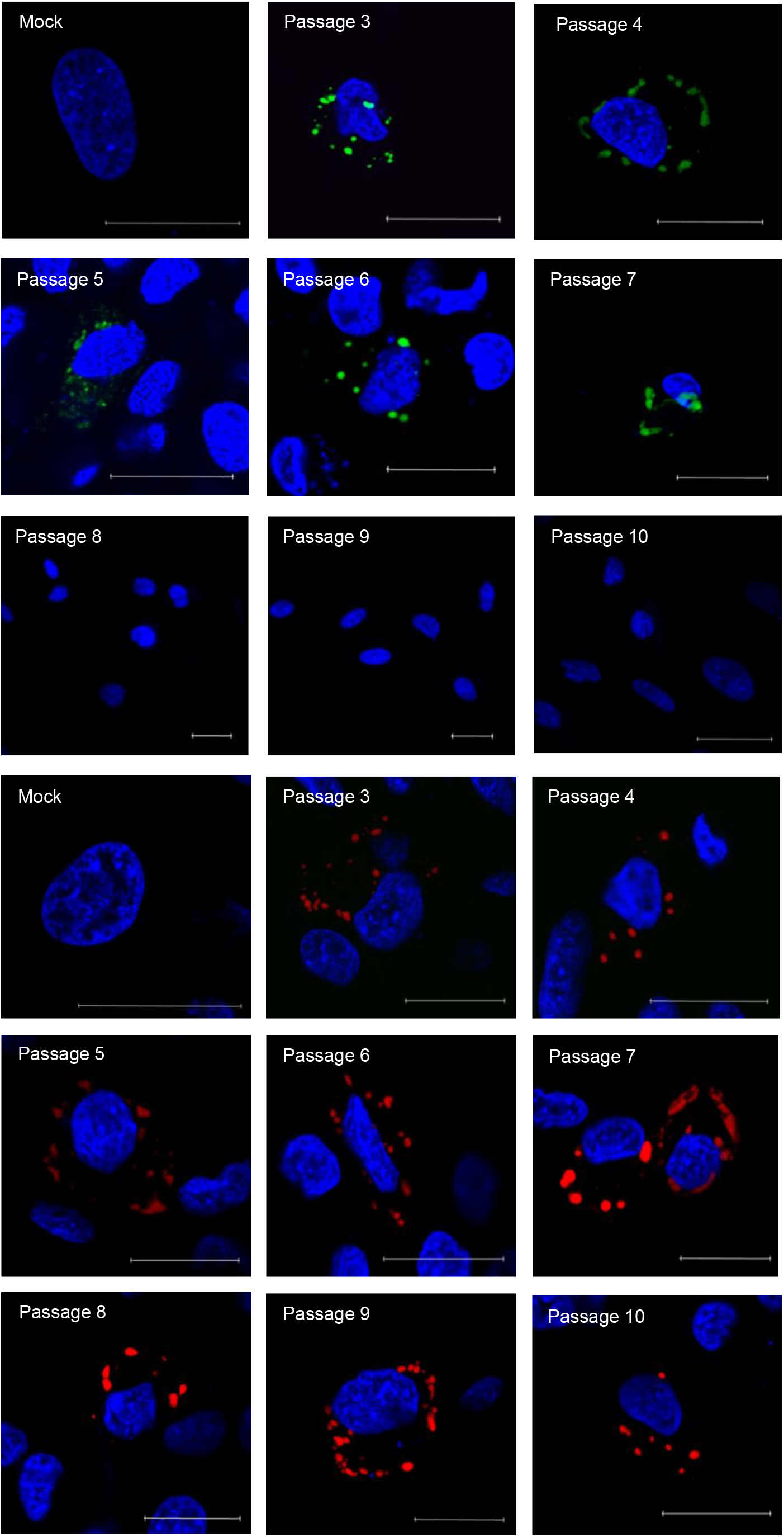
The PBG98-VP1-GFP11 virus is stable for 7 passages and the PBG98-VP1-TC virus is stable for at least 10 passages. DF1 cells were infected with passage 3 of either the PBG98-VP1-GFP11 virus or the PBG98-VP1-TC virus. A sample of the supernatant was obtained 24 hours post infection (hpi) and passaged onto additional DF1 cells. This was repeated for a total of 10 passages for each virus. Supernatant samples from each passage of the PBG98-VP1-GFP11 virus were added to DF-1 cells that had previously been transfected with the GFP1-10 plasmid, and fixed, stained with DAPI, and imaged 20 hpi. The supernatant samples from each passage of the PBG98-VP1-TC virus was added to DF-1 cells that were fixed and stained with ReAsH and DAPI, and imaged 20 hpi. Scale bars are 20μm in length.

**Fig. S2.**
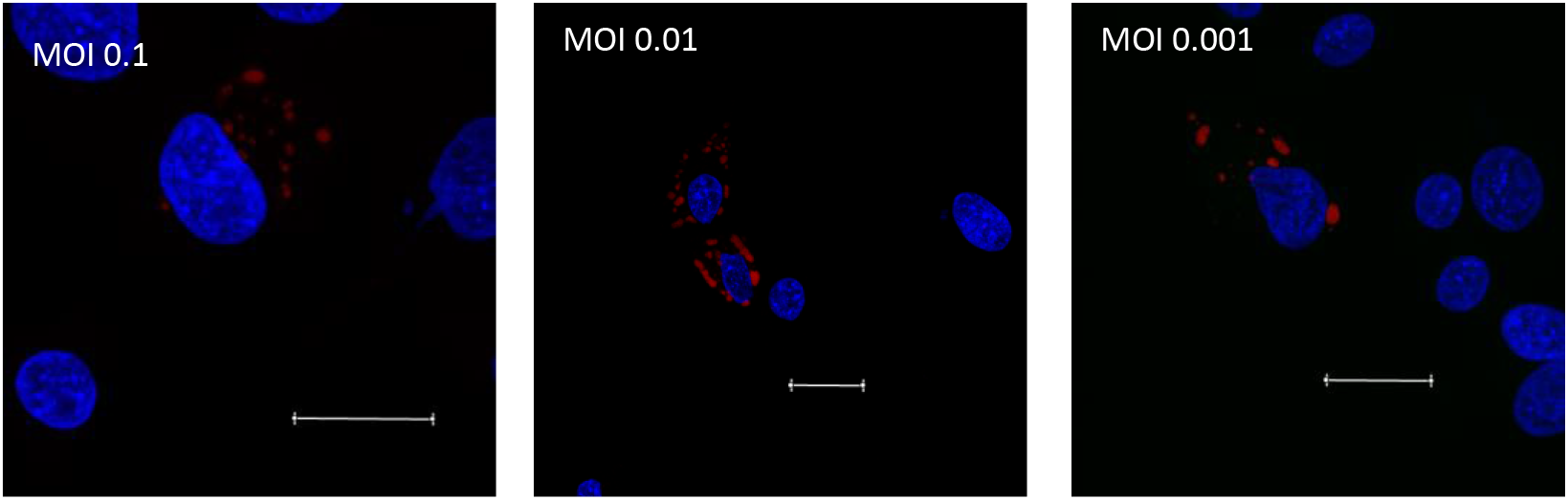
multiple VFs are observed per infected cell, irrespective of MOI. DF1 cells were infected with the PBG98-VP1-TC virus at an MOI of 0.1, 0.01 or 0.001. Cells were fixed at 20 hours post-infection and stained with ReAsH and DAPI. Scale bars are 20μm in length.

**Movie S1. VFs coalesce in the cytoplasm of infected cells between 16 and 18 hours post infection**. DF1 cells were transfected with a plasmid expressing GFP1-10 and infected 24 hours post-transfection with the PBG98-VP1-GFP11 virus at an MOI of 1. One live infected cell was imaged from 16-18 hours post infection; one image obtained every 4 minutes.

**Movie S2. VFs coalesce in the cytoplasm of infected cells between 22 and 24 hours post infection**. DF1 cells were transfected with a plasmid expressing GFP1-10 and infected 24 hours post-transfection with the PBG98-VP1-GFP11 virus at an MOI of 1. One live infected cell was imaged from 22-24 hours post infection; one image obtained every 3 minutes.

**Movie S3. VF coalescence is dependent on an intact microtubule network**. DF1 cells were transfected with a plasmid expressing GFP1-10 and infected 24 hours post-transfection with the PBG98-VP1-GFP11 virus at an MOI of 1. Two hours post-infection, cells were treated with nocodazole. One live infected cell was imaged from 20-22 hours post infection; one image obtained every 3 minutes.

**Movie S4. VF coalescence is dependent on an intact actin cytoskeleton**. DF1 cells were transfected with a plasmid expressing GFP1-10 and infected 24 hours post-transfection with the PBG98-VP1-GFP11 virus at an MOI of 1. Two hours post-infection, cells were treated with cytochalasin-D. One live infected cell was imaged from 20-22 hours post infection; one image obtained every 3 minutes.

